# Alternative Splicing Directs PMCA2 to Lysosomes and is Linked to Neurodegeneration

**DOI:** 10.1101/2025.10.01.679724

**Authors:** M.E. Fernández-Suárez, R. Bush, J.W. Brenton, G. Pereira, M. Grant-Peters, R. Reynolds, D. te Vruchte, D. Shepherd, Y. Weng, E. Artaza-Fernández, P. Lis, L. Sanchez-Pulido, A. Morgan, L. Davis, D. Gómez-Coronado, E. Eden, D. Alessi, C.P. Ponting, A. Galione, S. Newstead, J. Hardy, S. Patel, M. Ryten, F.M. Platt

## Abstract

Plasma membrane calcium ATPases (PMCAs) are believed to function exclusively at the plasma membrane where they expel calcium from the cytosol. We have unexpectedly identified a splice variant-dependent localisation of the PMCA isoform PMCA2 to the lysosome, where it forms an evolutionarily conserved complex with NPC1, the lysosomal membrane protein defective in the rare lysosomal storage disease Niemann-Pick disease type C (NPC). This interaction is required for lysosomal Ca^2+^ homeostasis and implicates PMCA2 as a mediator of Ca^2+^ uptake into lysosomes. Disruption of the NPC1-PMCA2 complex contributes to the pathophysiology of both Niemann-Pick disease type C and Parkinson’s disease, revealing an unrecognised intracellular function for PMCA2 and a shared mechanism linking lysosomal Ca^2+^ and lipid regulation in neurodegeneration.

## Introduction

Precise regulation of intracellular calcium (Ca^2+^) homeostasis is essential for neuronal function and survival. Plasma membrane calcium ATPases (PMCAs) are high-affinity Ca^2+^ pumps that transport cytosolic Ca^2+^ across the plasma membrane to the extracellular environment (*1*). Of the four PMCA isoforms in humans (PMCA1–4), PMCA2 and PMCA3 are highly expressed in neurons in the brain, with PMCA2 particularly enriched in cerebellar Purkinje cells (*1*). Loss-of-function mutations in PMCA2 or PMCA3 cause progressive cerebellar ataxia (*2–4*), and reduced PMCA2 expression has been associated with increased vulnerability of dopaminergic neurons in Parkinson’s disease (PD) (*5*). Moreover, soluble α-synuclein, implicated in PD pathogenesis, directly enhances PMCA1/2 activity, underscoring the relevance of PMCA regulation in neurodegeneration (*6*). To date, all functional roles of PMCAs have been attributed to its localisation at the plasma membrane. However, the genes encoding the PMCAs are alternatively spliced, raising the possibility that they could have additional localisations and functions.

In a yeast screen, we previously identified PMC1, the single yeast orthologue of mammalian PMCAs, as a direct interactor of Ncr1, the yeast orthologue of mammalian NPC1 (*7*). This interaction was conserved in mammalian cells (*7*). NPC1 is a thirteen-transmembrane lysosomal protein encoded by *NPC1* that is mutated in the rare neurodegenerative lysosomal storage disorder, Niemann-Pick disease type C (NPC). NPC1 deficiency results in disrupted lysosomal lipid homeostasis, impaired lysosome: ER membrane contact site formation and, critically, lowered levels of lysosomal luminal Ca^2+^ (*8*), potentially due to a Ca^2+^ store filling defect, a feature also reported in PD (*9*). However, the molecular mechanism underlying Ca^2+^ import into the lysosome remains incompletely understood.

Here, we show that alternative splicing directs specific PMCA2 isoforms to the lysosome, where they form a complex with NPC1 to regulate lysosomal Ca^2+^ levels. These findings have uncovered a conserved calcium regulatory axis with broad implications for lysosome function and neurodegenerative diseases. This study also highlights that splice variants can differentially affect subcellular organelle targeting of proteins, expanding the function of proteins beyond their canonical roles.

## Results

### PMCA2 colocalises with NPC1 in the lysosome

We previously reported that NPC1 and PMCA2 from rat cerebellum co-immunoprecipitate, indicating a direct interaction between these proteins (*7*). To determine whether this interaction occurs at lysosomes, we examined endogenous PMCA2 localisation in rat PC12 cells that can be differentiated into neurons, which express PMCA2 (*10, 11*). Confocal microscopy using a pan-PMCA2 antibody revealed robust lysosomal colocalization of PMCA2 and NPC1, with no detectable plasma membrane signal of PMCA2 (Fig. 1a). Staining controls confirmed antibody specificity and no bleed-through between channels (Supplementary Fig. 1). To further validate PMCA2 detection, we generated stable *Atp2b2* knockdown (KD) PC12 cells. The punctate lysosomal pattern observed in WT cells was absent in KD cells, supporting the antibody specificity and lysosomal localisation of PMCA2 (Supplementary Fig. 2).

**Fig. 1.**
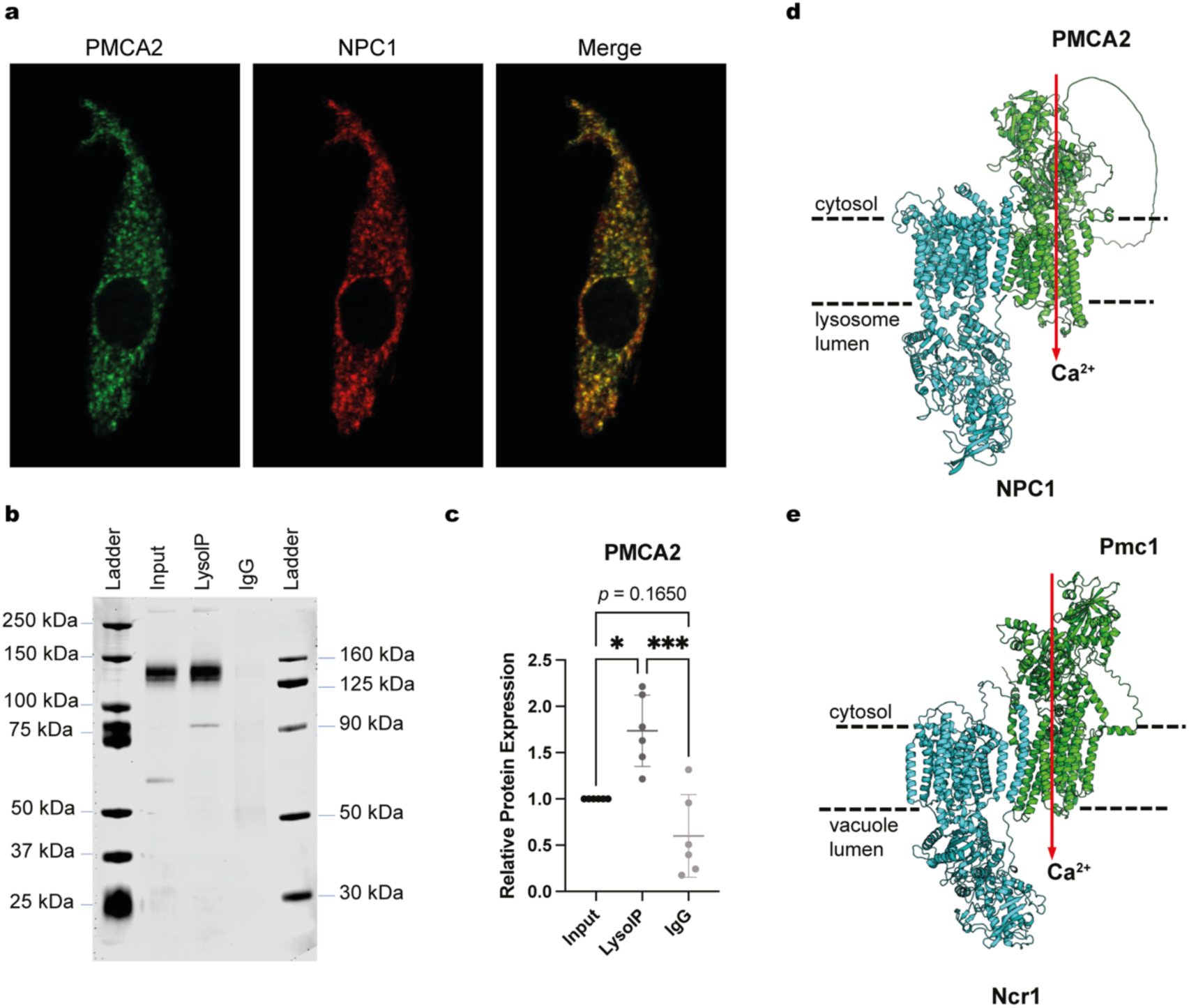
PMCA2 is present in the lysosome. (a) Immunocytochemistry and confocal microscopy showed that endogenous PMCA2 colocalises with NPC1 in the lysosome of PC12 cells differentiated into neurons. NPC1 and PMCA2 were detected using primary antibodies directly labelled with CoraLite-555 and CoraLite-488, respectively. Representative image of four independent experiments, Pearson’s coefficient r = 0.8527 ± 0.0415 standard deviation corresponding to the mean of four independent experiments, calculated using the mean of r from 10 fields per experiment. (b) Representative image of Tagless LysoIP followed by Western blot probing for PMCA2 in mouse cerebellar tissue, *n* = 6 mice. (c) Quantification of PMCA2 enrichment in lysosomal fraction from LysoIP experiments, *n* = 6 mice, lines represent the mean and standard deviation, one-way Anova test, \**p* < 0.05, \*\*\**p* < 0.001. (d) AlphaFold 3 model of the PMCA2-NPC1 complex in the lysosomal membrane. Directionality of Ca^2+^ transport is represented with a red arrow (e) AlphaFold 3 model of yeast orthologues Ncr1 and Pmc1 in the vacuolar membrane exhibiting a similar orientation and interaction.

To independently confirm this compartmentalisation, we isolated lysosomes from WT mouse cerebellum using Tagless LysoIP (*12*). Western blotting detected a ∼130 kDa PMCA2 band enriched in the lysosomal fraction relative to an isotype control (Fig. 1b and c), consistent with endogenous PMCA2 being localised at the lysosome.

To explore how PMCA2 may interact with NPC1, we modelled the complex using AlphaFold 3. The predicted structure positioned both the N- and C-terminal domains of PMCA2 on the cytosolic side of the lysosomal membrane, compatible with a role in transporting Ca^2+^ from the cytosol into the lysosomal lumen (Fig. 1d). Strikingly, modelling of the yeast Pmc1/Ncr1 complex generated a similar topology (Fig. 1e), suggesting that the organisation and function of this complex are conserved across evolution.

### PMCA2 depletion disrupts lysosomal Ca^2+^ signalling

The lysosomal localisation and predicted orientation of PMCA2 suggested that it could contribute to lysosomal Ca^2+^ filling. To test this hypothesis, we used *Atp2b2* KD PC12 cells differentiated into neurons and loaded them with the cytosolic Ca^2+^ indicator Fura-2. Lysosomal Ca^2+^ release was evoked with the selective TPC2 agonists TPC2-A1N and TPC2-A1P (*13*). KD neurons exhibited significantly reduced agonist-stimulated Ca^2+^ release compared with control neurons (Fig. 2a-c), indicating a deficit in lysosomal Ca^2+^ stores when PMCA2 expression is reduced.

**Fig. 2.**
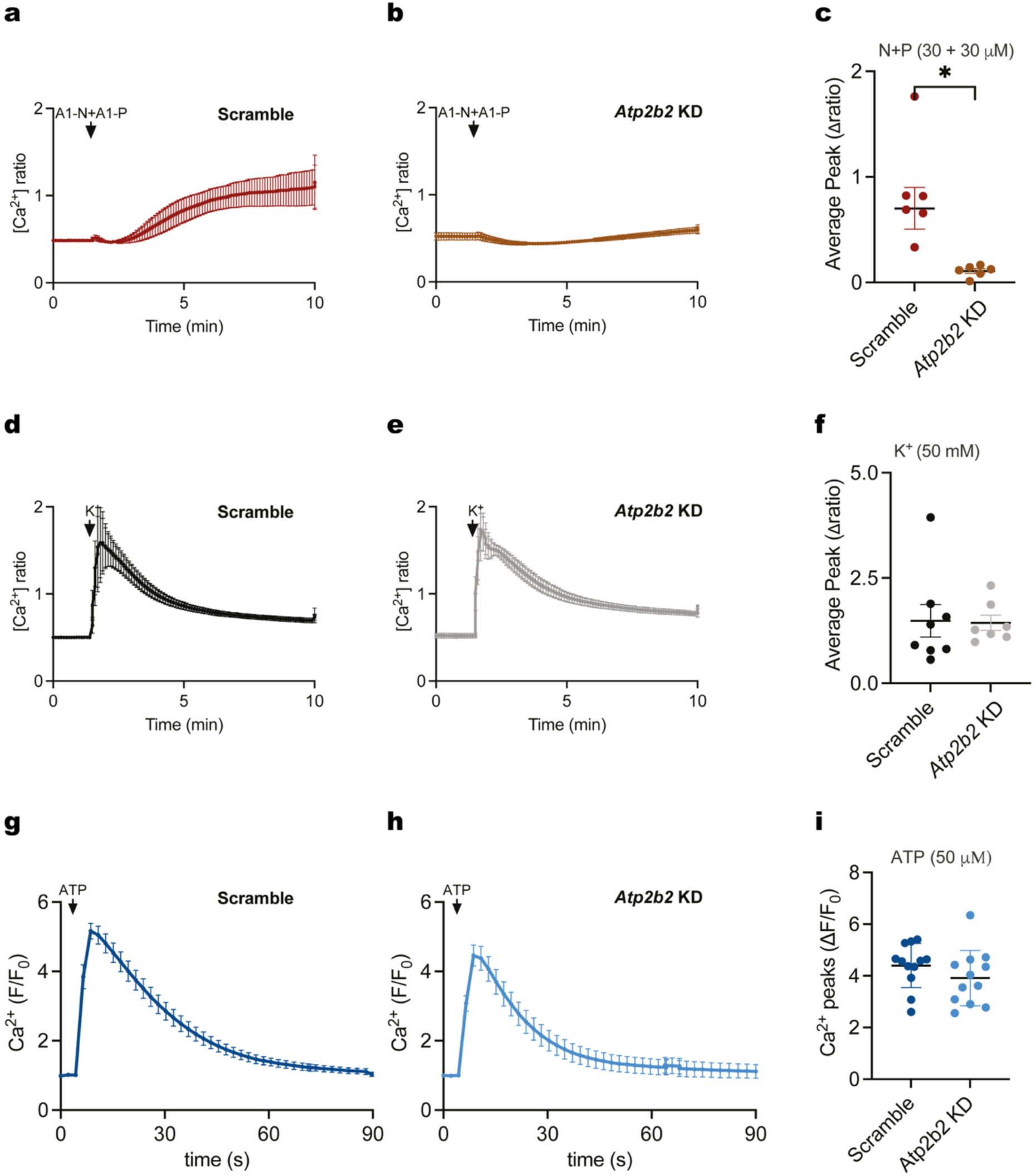
Lysosomal Ca^2+^ content is reduced in *Atp2b2* KD cells. PC12 cells stably expressing scramble or *Atp2b2* targeting shRNA were preloaded with the Ca^2+^ probe Fura-2. Ca^2+^ signal in response to a combination of the TPC2 agonists TPC2A1N (30 μM) and TPC2A1P (30 μM) in scramble (a) and *Atp2b2* KD cells (b). Summary data (c) quantifying the peak change in Ca^2+^ signal in scramble and *Atp2b2* KD cells stimulated with the TPC2 agonists. Each point represents the mean response from 30 cells per coverslip from three independent experiments, two coverslips per experiment. Ca^2+^ signal when scramble (d) and *Atp2b2* KD cells (e) were depolarised with KCl (50 mM). Summary data (f) quantifying the peak change in the Ca^2+^ signal in scramble and *Atp2b2* KD cells depolarised with KCl. Each point represents the mean response from 30 cells per coverslip from four independent experiments, two coverslips per experiment. Ca^2+^ signal when scramble (g) and *Atp2b2* KD cells (h) preloaded with Calbryte-590 were stimulated, in Ca^2+^-free medium, with 50 µM ATP. Summary data (i) quantifying the peak change in the Ca^2+^ signal in response to ATP. Each point represents the mean response from > 100 cells per well from four independent experiments, two to four wells per experiment. Lines represent the mean and standard deviation, \**p* < 0.05 calculated by unpaired t-test.

Basal cytosolic Ca^2+^ levels were unchanged between groups (Supplementary Fig. 3a), suggesting that global Ca^2+^ homeostasis was preserved. To confirm that the differentiation state and general excitability of both groups were intact, neurons were stimulated with 50 mM KCl to depolarize voltage-gated Ca^2+^ channels. Both control and KD cells responded with robust Ca^2+^ influx (Fig. 2d–f), demonstrating that the KD phenotype is specific to lysosomal Ca^2+^ handling. GFP expression from the shRNA cassette did not interfere with Fura-2 imaging (Supplementary Fig. 3b-d).

Although the inhibition of TPC2-dependent Ca^2+^ responses in KD neurons is consistent with a reduction in lysosomal Ca^2+^ filling, an alternative explanation needed to be excluded. It is well known that the small Ca^2+^ release from lysosomes is amplified by the large, secondary recruitment of ER Ca^2+^-release channels such as IP_3_Rs via the process of Ca^2+^-induced Ca^2+^ release (CICR) (*14*); hence, Fura-2 signals were potentially dominated by the large secondary ER component. It was therefore conceivable that the reduced responses to TPC2 agonists in KD neurons reflected impaired ER Ca^2+^ release rather than a primary lysosomal defect.

To address this possibility, we directly assessed ER Ca^2+^ release by stimulating purinergic receptors with ATP, which activates IP₃-mediated Ca^2+^ release from the ER (*14*). Cytosolic Ca^2+^ signals were monitored using the low-affinity red indicator Calbryte-590 (Kd ≈ 1.2 µM), which minimises saturation at peak Ca^2+^ levels relative to Fura-2 (Kd ≈ 0.22 µM). Experiments were performed in Ca^2+^-free medium to monitor Ca^2+^ release from intracellular stores only. The peak Ca^2+^ amplitudes of ATP responses were not significantly different between Scrambled and KD neurons (Fig. 2g-i). Thus, ER-dependent amplification cannot account for the reduced responses to TPC2 agonists when PMCA2 is defective, strengthening the argument for an effect upon the lysosomes themselves.

Collectively, these findings position PMCA2 as a previously unrecognised contributor to lysosomal Ca^2+^ filling and reveal that PMCA2 loss disrupts lysosomal Ca^2+^ signalling.

### Several PMCA splice variants contain a lysosomal targeting motif

The PMCA gene family (ATP2B1-4) generates multiple isoforms through alternative splicing at two major sites, splice site A and C (*1*). We found that variation at site C can incorporate a canonical tyrosine-based lysosomal targeting motif, YEGL (*15*). Analysis of human and rat transcripts revealed that several isoforms, PMCA1b, 1c, 1d; PMCA2b; and PMCA3b, 3c, 3d, include the YEGL motif (Fig. 3a; Supplementary Fig. 4). No PMCA4 isoform examined contained this motif, suggesting isoform-specific differences in intracellular trafficking.

**Fig. 3.**
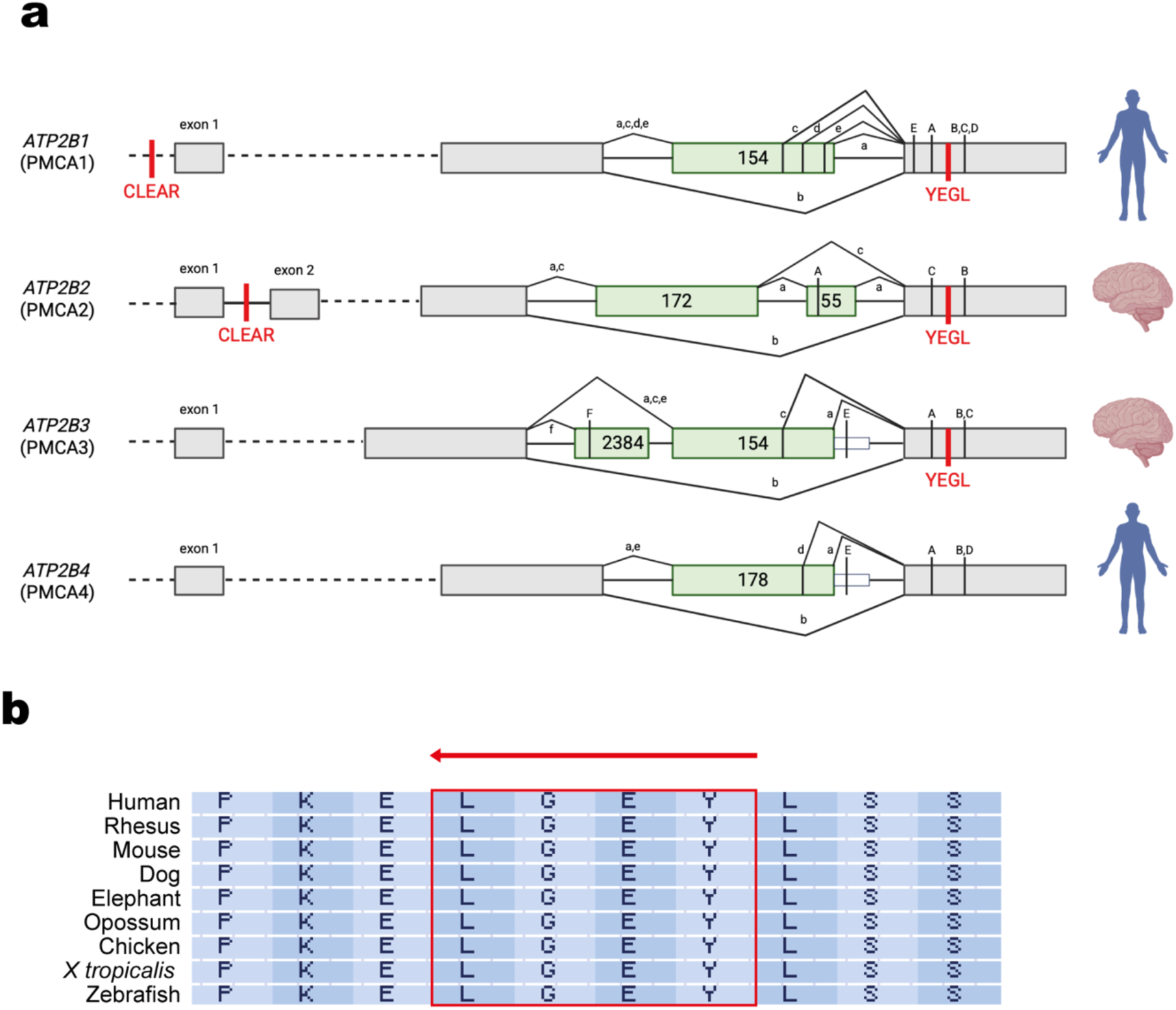
Specific PMCA splice variants contain the lysosomal targeting motif YEGL. *ATP2B1* and *ATP2B2* contain CLEAR sequences. (a) PMCA site C alternative splicing in human can include or exclude the lysosomal targeting motif YEGL. The exon structure of the different regions affected by alternative splicing is shown for each of the four PMCA genes. Constitutively spliced exons are indicated as grey boxes. Alternatively-inserted exons are shown in green. Lowercase letters represent resulting splice variants. Capital letters indicate the position of the translated stop codons. In PMCA3 and PMCA4, splice variant “e” results from a read-through of the exon into the following intron (indicated as small white boxes). The sizes of alternatively spliced exons are given as nucleotide numbers. Splice variants shown have supporting sequence data in the literature, RefSeq or Ensembl at the time of submission of this work. (b) Example of the YEGL motif alignment in the *ATP2B2* gene from different species.

We also identified CLEAR elements, TFEB response sequences that regulate lysosomal gene expression (*16*), in ATP2B1 and ATP2B2 in both human and rat genomes (Supplementary Fig. 4). The presence of CLEAR sequences indicates that PMCA1 and PMCA2 transcription may be coordinated with lysosomal biogenesis and function.

Next, we assessed the evolutionary conservation of the lysosomal targeting YEGL in *ATP2B1* and *ATP2B2* genes, using 100 vertebrate genomes aligned by UCSC. All but one (Lamprey) *ATP2B1* ortholog and all but two (duck-billed platypus and Lamprey) *ATP2B2* orthologs retained YEGL (Fig. 3b), underscoring the motif’s strong evolutionary preservation. The species that did not contain this motif had variations on this sequence that require testing to see if they do direct these proteins to the lysosome. Yeast PMC1 carries a similar tyrosine-based vacuolar targeting motif (YYFL), consistent with PMCAs originally functioning in acidic organelles before later diversification (*17, 18*).

To experimentally assess isoform-specific localisation, we generated PC12 cells expressing GFP-tagged PMCA2 splice variants under a tetracycline-inducible promoter. As expected, PMCA2wa localised primarily to the plasma membrane. YEGL-containing variants exhibited mixed localisation, with strong plasma membrane expression but also intracellular signals when overexpressed (Supplementary Fig. 5). This differed from the predominantly lysosomal localisation of endogenous PMCA2, suggesting that correct trafficking may require regulated expression, stoichiometric complementarity with NPC1 levels or accessory factors.

### PMCA2 deficiency is a key component of the NPC disease pathway

We previously reported that expression of the *Atp2b2* (PMCA2) gene is reduced in the cerebellum of *Npc1^-/-^* mice, an NPC model that recapitulates the biochemical and phenotypic features of the human disease (*7*). Here, analysis of individual splice variants revealed PMCA2b as the predominant isoform in the cerebellum of wild-type mice (Fig. 4a), followed by PMCA2a. All isoforms were significantly downregulated in *Npc1^-/-^* cerebellum, yielding an overall 67% reduction in *Atp2b2* expression (Fig. 4b). Because PMCA2 contributes to lysosomal Ca^2+^ filling, its reduction in NPC may explain the low lysosomal Ca^2+^ levels previously reported in this disorder.

**Fig. 4.**
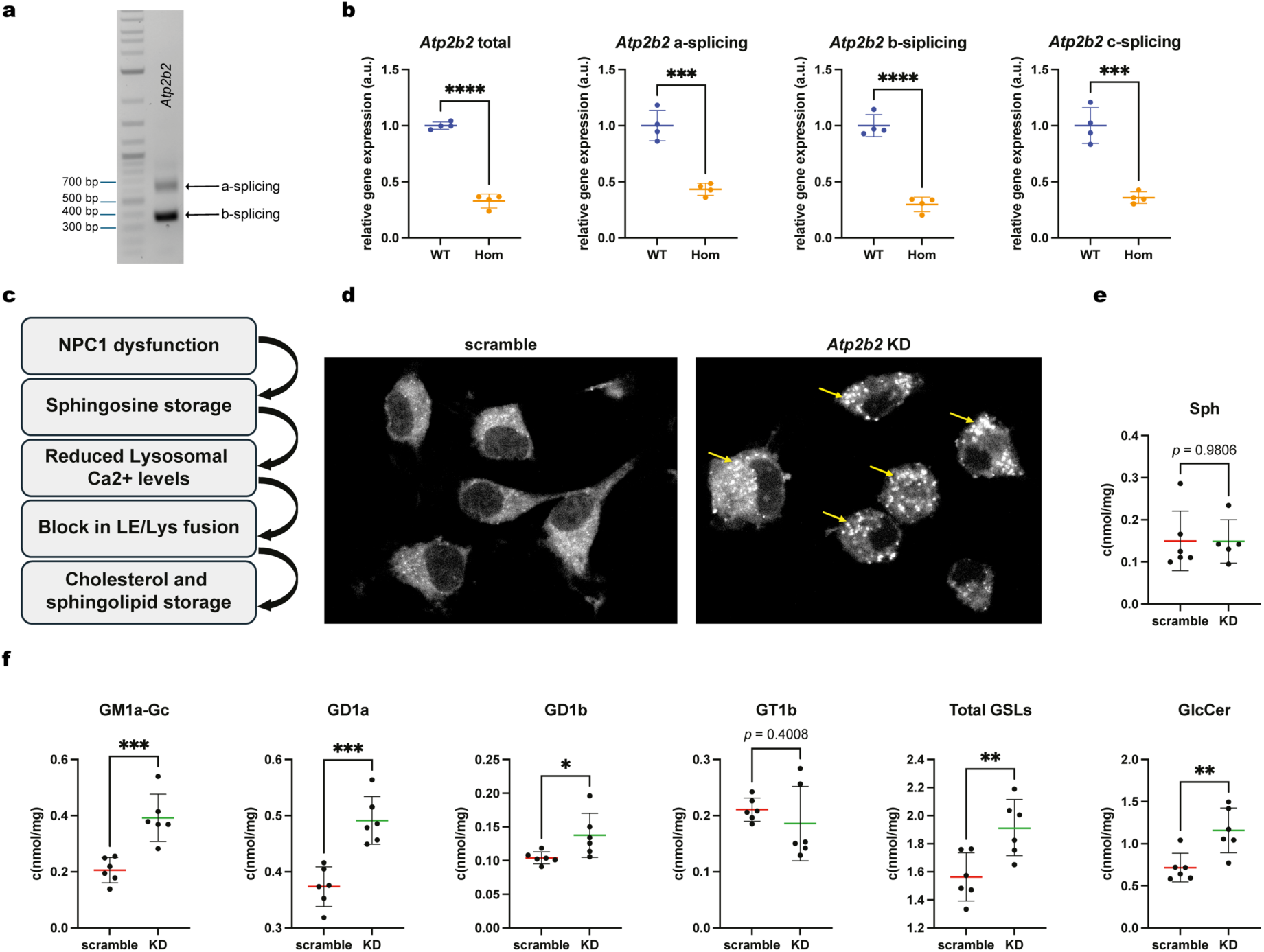
*Atp2b2* splice variants are reduced in NPC and, reciprocally, *Atp2b2* deficiency induces multiple NPC phenotypes, but not sphingosine accumulation. (a) Gel electrophoresis of PCR products showing the relative abundance of the different *Atp2b2* splice variants in the cerebellum of 8-week-old WT mice. (b) Relative gene expression measured by qPCR of different splice variants of *Atp2b2* in WT and *Npc1^-/-^* (Hom) cerebellums of 8-week-old *Npc1^-/-^* mice, *n* = 4 mice. (c) Diagram of the proposed NPC pathogenic cascade (*8*). (d) Filipin staining of free cholesterol in scramble and *Atp2b2* KD cells, representative image of 3 independent experiments. Sphingosine (e) and GSLs (f) concentrations in scramble vs *Atp2b2* KD cells, (*n* = 6 scramble and 5 KD for sphingosine, due to the exclusion of an outlier detected using Grubbs’ test; *n* = 6 for GSLs). Lines represent the mean and standard deviation, \**p* < 0.05, \*\**p* < 0.01, \*\*\**p* < 0.001, \*\*\*\**p* < 0.0001 calculated by unpaired t test.

We next investigated whether *Atp2b2* KD PC12 neurons exhibit cellular phenotypes associated with NPC (Fig. 4c). Control neurons showed diffuse cholesterol staining visualised with filipin, whereas KD neurons accumulated punctate intracellular cholesterol, consistent with lysosomal cholesterol accumulation (Fig. 4d). Sphingosine levels were unchanged (Fig. 4e). However, several glycosphingolipids associated with NPC pathology, including GM1a-Gc, GD1a, GD1b, and GlcCer, were elevated (Fig. 4f).

These results demonstrate that PMCA2 reduction alone reproduces multiple canonical NPC phenotypes, indicating that reduced PMCA2 expression contributes substantially to the disease pathway and may act upstream of several downstream metabolic defects.

Many genes causative of lysosomal storage diseases, including *NPC1*, have been found in the heterozygous state to be over-represented in Parkinson’s disease (*19, 20*). We therefore investigated whether mRNA levels of *NPC1* and its interacting partners (including PMCA2) were altered in Lewy body diseases.

### The splicing of PMCAs is altered in Lewy body diseases

Lewy body diseases (LBD), including Parkinson’s disease (PD), Parkinson’s disease with dementia (PDD), and dementia with Lewy bodies (DLB), exhibit widespread lysosomal dysfunction. To investigate whether PMCA dysregulation contributes to LBD pathology, we analysed single-nucleus and bulk RNA-seq datasets from human anterior cingulate cortex tissue across 28 individuals (controls, PD, PDD, DLB; n = 7 per group) (*21*).

Single-nucleus analysis revealed significant reductions in *ATP2B2* expression across several cell types in PD, PDD and DLB, with the strongest and most consistent effects in excitatory neurons. Multiple glial cell types, including oligodendrocyte precursor cells, oligodendrocytes, and microglia, also showed significant downregulation in at least one disease state (Fig. 5a).

**Fig. 5.**
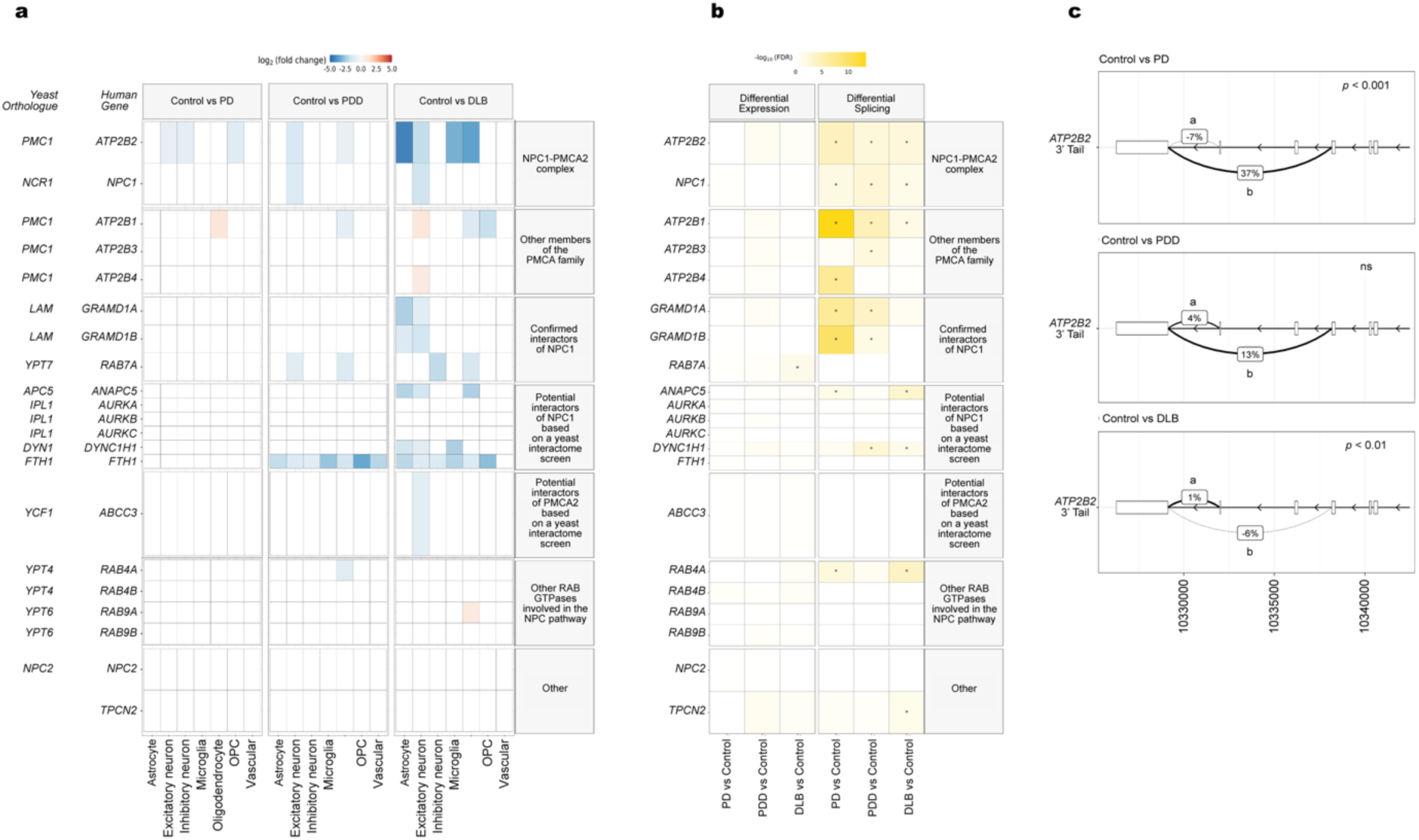
PMCA2, NPC1 and their wider interactome gene expression and splicing are altered in Lewy body diseases. (a) Gene expression changes using single-nucleus RNA sequencing in post-mortem human brain samples from PD, PDD and DLB cases and controls. Fill of the tile indicates the significant (FDR < 0.05) log2(fold change), with non-significant results (FDR > 0.05) coloured white. (b) Gene expression and differential splicing of *NPC1* and potential NPC1 interacting partners using bulk RNA sequencing in PD, PDD and DLB compared to controls. The fill colour of each tile represents the −log10(FDR) for the gene (left panel) or the most significant spliced junction cluster within each gene (right panel), with FDR significance <0.05 indicated by an *. (c) *ATP2B2* splice variants demonstrate differential splicing profiles in Lewy body disorders. Sashimi plots showing delta percent spliced in comparison to controls for junctions producing alternative *ATP2B2* splice variants a and b. (a) *n* = 7 patients per group. (b and c) *n* = 5 controls, *n* = 7 PD, *n* = 6 PDD and *n* = 6 DLB. PD: Parkinson’s disease, PDD: PD with dementia, DLB: dementia with Lewy bodies.

We expanded the analysis to NPC1 and 20 additional genes encoding confirmed or proposed NPC1 interacting partners (*7, 22–24*). Of these, 13 were significantly differentially expressed (the majority downregulated) in at least one cell type or disease group (Fig. 5a). Notably, FTH1, a gene central to iron metabolism, showed broad downregulation across multiple LBD cell types.

Given the importance of alternative splicing for PMCA localisation, we next examined differential transcript usage in high-depth bulk RNA-seq data. Ten of the 21 genes analysed, including *ATP2B1*, *ATP2B2*, and *NPC1*, were significantly differentially spliced in PD, PDD, or DLB (Fig. 5b). Because inclusion of the YEGL motif determines PMCA lysosomal targeting, we focused on *ATP2B2* splicing events affecting this region. Variant-b junction usage, which introduces the YEGL motif, was significantly increased in PD (ΔPSI = +37%, FDR < 0.001) and decreased in DLB (ΔPSI = –6%, FDR < 0.01) (Fig. 5c). Similar patterns in *ATP2B1* and *ATP2B4* (Supplementary Table 1) point to widespread disruption of PMCA lysosomal targeting in LBD.

Together, these findings indicate that PMCA expression and splicing are perturbed not only in NPC but also across LBDs, suggesting a shared vulnerability to altered lysosomal Ca^2+^ handling pathways in neurodegenerative diseases more broadly.

## Discussion

In this study, we surprisingly found that PMCA2 localises to the lysosome. A possible explanation for this unexpected localisation of PMCA is the presence of a canonical tyrosine-based lysosomal targeting motif (YEGL), which we identified in specific splice variants of PMCA genes. In addition, both PMCA1 and PMCA2 harbour CLEAR (Coordinated Lysosomal Expression and Regulation) elements, further supporting a lysosomal role for these isoforms. Notably, the YEGL motif was conserved across 98% of the 100 species examined, suggesting it plays a fundamental and evolutionarily conserved role. Intriguingly, orthologues of PMCAs in yeast and *Dictyostelium* localise to the vacuole, the equivalent of the lysosome in these organisms, raising the possibility that plasma membrane targeting of PMCAs may have evolved secondarily to an ancestral lysosomal localisation and function.

Interestingly, when PMCA2 splice variants containing the lysosomal targeting motif were overexpressed, they localised not only to intracellular compartments but also to the plasma membrane. This dual localisation suggests that intracellular retention may require interaction with specific partner proteins in the lysosome. In the absence of sufficient binding partners, may, by default, insert in the plasma membrane, raising the possibility that stoichiometric balance with interacting partners such as NPC1 also affects its subcellular distribution.

Our results are consistent with the hypothesis that the function of PMCA2 at the lysosomal membrane is to complex with NPC1, which tethers the lysosome to the ER (*22*) and that, then, PMCA2 fills the lysosome with Ca^2+^ at lysosome: ER contact sites. The lysosome is a regulated Ca^2+^ store and signalling hub. Lysosomal Ca^2+^ release is needed for vesicular trafficking, endosome/lysosome fusion, and phagocytosis (*25*). Several proteins have been described to facilitate calcium release from the lysosome, such as the TPC2 and TRPML1 channels (*25*) but the mechanisms for lysosomal Ca^2+^ filling remain incompletely understood (*26*). TMEM165 has recently emerged as a candidate (*27, 28*). However, it has long been speculated that both passive and active Ca^2+^ transporters would be required (*14*). Our study presents PMCA2 as an additional candidate for lysosomal Ca^2+^ filling, specifically at NPC1-mediated contact sites between the lysosome and ER.

After demonstrating its lysosomal localisation and interaction with NPC1 (*7*), we modelled its orientation in the limiting membrane of the lysosome. Our results using AlphaFold favoured an orientation compatible with transporting calcium into the lysosome. Consistent with this, when PMCA2 was stably knocked down, the release of Ca^2+^ from the lysosome induced by the TPC2 agonists TPC2-A1-N and TPC2-A1-P was significantly reduced compared to control cells. These results are compatible with lower levels of lysosomal Ca^2+^ when PMCA2 is defective, supporting a role for PMCA2 in mediating lysosomal Ca^2+^ filling. However, our measurements were indirect, relying on monitoring cytosolic Ca^2+^ upon opening of a lysosomal ion channel as opposed to direct measurements of Ca^2+^ within the lysosomal lumen. The latter is challenging due to the acidic interior of lysosomes. Nevertheless, lysosomal calcium filling by PMCA2 clearly merits further investigation.

When NPC1 is defective, the first event in the pathogenic cascade is the accumulation of sphingosine that leads to the reduction of lysosomal Ca^2+^ levels (*8*). The fact that in cells with functional NPC1, knockdown of PMCA2 does not result in the accumulation of sphingosine is consistent with NPC1-dependent sphingosine transport being upstream of the lysosomal Ca^2+^ defect. Our discovery that PMCA2 expression is reduced in *Npc1*-null mice links this Ca^2+^ transporter to the NPC pathway, explaining for the first time why lysosomal Ca^2+^ levels are reduced when NPC1 is defective. This hypothesis is further supported by the fact that when we knockdown PMCA2, lysosomal Ca^2+^ levels are also reduced. Another link between PMCA2 and the NPC pathway is that in cells with intact NPC1 but reduced levels of PMCA2, lipids such as cholesterol and the majority of GSLs accumulate, as observed in NPC cells. These data suggest that the NPC1/PMCA axis is a hub regulating lysosomal Ca^2+^ and lipid homeostasis at NPC1 mediated contact sites between the lysosome and ER.

It is interesting that mutations in PMCA2 in humans and mouse models lead to ataxia (*2, 3*), another characteristic phenotype of NPC. Furthermore, PMCA2 is highly expressed in the cerebellum, particularly in Purkinje neurons, a cell type that is uniquely susceptible to degeneration in NPC (*29*).

Importantly, the reduction in PMCA2 gene expression was not limited to NPC but was also observed in several cell types in PD, PDD, and DLB. This is highly relevant as it has been reported that a reduction in *ATP2B2* (PMCA2) increases neuronal vulnerability in *in vitro* PD models (*5*). In this study, we found that a reduction in PMCA2 leads to GSL dyshomeostasis, which has also been reported in PD (*30, 31*). Furthermore, low lysosomal Ca^2+^ levels have also been reported in PD (*9, 26*). Since mutations in NPC1 are overrepresented in PD (*19, 20*), taken together with our findings in this study, we hypothesise that the NPC1-PMCA2 complex is part of a wider interactome in the lysosome whose dysregulation plays a key role in NPC and Lewy body pathologies. Supporting this hypothesis, gene expression of *NPC1, ATP2B2* and several of their confirmed and potential binding partners are also dysregulated in different brain cell types in Lewy body diseases.

A key finding in this study was a profound PD-related change in splice variant usage in several members of the wider NPC1 interactome. Changes in the splice variant expressed could, potentially, change the protein localisation, regulation or function in the disease state. This result highlights the need for more comprehensive studies that include not only total expression but also differential splice variant usage.

In conclusion, our study uncovers for the first time a previously unrecognised, differential splicing-dependent role for PMCA2 involved in lysosomal Ca^2+^ filling, expanding the known functions of the PMCA family beyond its canonical activity at the plasma membrane. We show that alternative splicing directs specific PMCA2 isoforms to the limiting membrane of the lysosome, where they interact with NPC1 to restore luminal Ca^2+^ levels. This mechanism is essential for lysosomal function, and its disruption results in lysosomal Ca^2+^ depletion and lipid accumulation, features common to Niemann-Pick disease type C and Parkinson’s disease. These findings define a novel function for the NPC1–PMCA2 complex in lysosomal Ca^2+^ homeostasis and highlight how alternative splicing can diversify the functional landscape of well-characterised genes. More broadly, our work underscores the importance of isoform-specific regulation in organelle biology and opens new avenues for understanding and targeting lysosomal dysfunction in neurodegenerative diseases.

## Materials and Methods

### Materials

RPMI 1640 culture medium was obtained from Sigma. Foetal bovine serum (FBS), horse serum and Glutamax for supplementing the medium were obtained from Gibco. NGF-β from rat (N2513), and the antibiotics penicillin and streptomycin were obtained from Sigma.

### Low-density lipoprotein (LDL) and lipoprotein-deficient serum (LPDS) preparation

Human LDL (1.019–1.063 kg/l) was isolated by vertical rotor density gradient ultracentrifugation of plasma from hypercholesterolemic patients, as previously described (*32*). LPDS was prepared from FBS by ultracentrifugation at a density of 1.21 kg/l (*32*).

### Cells

PC12 cells (ATCC CRL-1721) were maintained in RPMI 1640 supplemented with 10% horse serum, 5% FBS, 2 mM L-alanyl-L-glutamine dipeptide in 0.85% NaCl (Glutamax), 100 U/ml penicillin and 100 U/ml streptomycin at 37 °C, in humidified atmosphere with 5% CO2. For differentiation to neurons, PC12 cells were plated onto poly-D-lysine-coated coverslips or flasks and incubated in RPMI 1640 supplemented with 1% horse serum, 100 ng/ml NGF-β and the abovementioned additives for four days. Medium was replaced on the second day.

For cholesterol distribution experiments, after differentiation, PC12 neurons were pre-treated with RPMI 1640 supplemented with 10% LPDS for 24 h. Subsequently, LDL (40 μg/ml cholesterol) was added to the culture. After 24 h, cells were fixed with 4% paraformaldehyde in PBS.

### Generation of stable *Atp2b2* KD and scramble PC12 cells

Lentiviral particles were prepared by transfecting HEK293T cells with Gag-Pol and pMDG packaging plasmids, and each of four rat *Atp2b2* or scrambled shRNA plasmid (TL709157, Origene). PC12 cells were transduced with the lentiviral constructs for 24 h. Then, we performed a kill curve with puromycin. After puromycin selection, *Atp2b2* expression was assessed by qPCR. Cells transduced with the shRNA leading to the greatest reduction (> 80%) of *Atp2b2* gene expression and scrambled shRNA cells were selected for further experiments.

### Animals

All experiments were conducted using protocols approved by the UK Home Office Animal Scientific Procedures Act, 1986. All animal work followed the ARRIVE guidelines. BALB/cNctr-Npc1^m1N^/J also named *Npc1^nih^*mice, were generated by heterozygote brother/sister matings obtained from Jackson Laboratory (Charles River, UK). Mice were bred and housed in individually ventilated cages (IVCs) under non-sterile conditions. Mice were culled at eight weeks of age with 800 mg/kg pentobarbital IP. For biochemical analysis, mice were transcardially perfused with ice-cold PBS and their brains removed and dissected.

### Microscopy

PC12 cells were plated onto poly-D-lysine-coated coverslips and differentiated into neurons over four days. Then cells were fixed with 4 % paraformaldehyde for 10 min and permeabilised with 0.1 % saponin in PBS for 1 hour at room temperature (RT). Cells were blocked with 10% horse serum in PBS for 1 hour. Primary polyclonal PMCA2 (PA1-915, Invitrogen) and monoclonal NPC1 (ab134113, abcam) antibodies were labelled with Flexable Coralite Plus-555 (KFA502, proteintech) and Flexable Coralite Plus-488 (KFA501, proteintech) for rabbit antibodies following the manufacturer’s instructions. Cells were probed with the conjugated antibodies overnight at 4°C, washed and mounted on slides using Prolong Gold mounting medium. Alternatively, after differentiation and incubation with LPDS and LDL as described above, PC12 neurons were fixed with 4% paraformaldehyde for 10 min and exposed to filipin (35 μg/ml in PBS) for 1 h. After washing, coverslips were mounted on slides with ProLong Gold. Confocal images were acquired using a Leica-SP8 confocal microscope with an LD 63X objective.

All image analysis were performed using ImageJ (FIJI) software (version 2.16.0/1.54g, ImageJ2). For quantification of colocalisation of NPC1 and PMCA2, the JACoP plugin (*33*) was used.

### Tagless LysoIP

Tagless LysoIP was performed as described by Saarela et al. (*12*) with minor modifications. Briefly, mouse cerebellum (whole cerebellum per data point) were collected and gently homogenised with 40 strokes of an electric homogeniser in 1 ml of potassium phosphate buffer saline (KPBS) supplemented with Halt protease inhibitor cocktail (Thermo Fisher Scientific) and PMFS (Cell Signaling Technology). Samples were centrifuged at 1000 g for 2 min at 4°C. Supernatant was collected, and 50 µl was reserved as input (whole tissue extract). The rest of the supernatant was divided into two tubes, each containing 100 µl of magnetic beads bound to antibodies against murine TMEM192 or isotype IgG. Magnetic beads were previously washed with KPBS. Supernatants were incubated with the magnetic beads for 5 min at 4°C on an orbital shaker. After incubation, the mixture was placed on a magnet for 1 min and the supernatant was removed. The beads were washed 3 times with ice cold KPBS supplemented with the protease inhibitors. Ripa buffer (Cell Signaling Technology) plus protease inhibitors was added to the beads and input. Samples with beads were placed on a magnet, and lysates were collected for western blot.

### Western blotting

Cells and lysoIP fractions were lysed in Ripa Buffer (Cell Signaling Technology) supplemented with a Halt^TM^ protease inhibitor cocktail (Thermo Fisher Scientific) and PMFS (Cell Signaling Technology). Protein expression was measured using the bicinchoninic acid method (BCA Protein Assay Kit, Pierce). Lysates were separated on 4–12% SDS–PAGE gels (Thermo Fisher Scientific) and then transferred to PVDF membranes. For Western blot normalisation, we used the Revert™ 700 Total Protein (LI-COR). After destaining, membranes were blocked with TBS-Intercept blocking buffer (LI-COR) and probed with primary polyclonal anti-PMCA2 (PA1-915, Invitrogen) antibody and secondary antibodies conjugated to IRDye 800CW (LI-COR). Fluorescence was detected using an Odyssey M scanner (LI-COR). Acquired images were visualised and quantified with the Empiria Studio Software (LI-COR).

### qPCR

Total RNA from mouse brain was isolated using the RNAeasy plus extraction kit (Qiagen) following the manufacturer’s protocol. cDNA was synthesised using the iScript cDNA synthesis kit (BIO-RAD). Real-time PCR amplification was performed on a LightCycler 480 II, using the PowerUp SYBER Green (Thermo Fisher Scientific), and specific primers (Supplementary table 2). Relative expression of each gene was determined by calculating the increase in CP (ΔCP) and normalising by the housekeeping gene Rpl37 (ribosomal protein L37).

### GSLs analysis

Glucosylceramide (GlcCer) and downstream GSLs were analysed as previously described (*34*) with minor modifications. Detailed protocols are available at protocols.IO for GlcCer and GSLs (*35*) determination. Briefly, GSLs were extracted from cell homogenates (200 μg of protein) in C:M 1:2 overnight. The mixture was centrifuged, chloroform and PBS were added to the supernatant and centrifuged. The lower phase was dried and resuspended in a small volume of C:M 1:3 and mixed with the upper phase. GSLs were recovered using C18 Isolute columns (100 mg, Biotage), and three quarters of the column elutant was dried and resuspended in ceramide glycanase buffer (50 mM sodium acetate pH 5.5, 1 mg/mL sodium taurodeoxycholate). rECGase 1 (prepared by Genscript) was added and samples incubated overnight. The other quarter of the column elutant was also dried and resuspended in ceramide glycanase buffer, but Cerezyme^®^ (Genzyme, Cambridge, MA) was added and incubated for 72 h. Released lipids were anthranilic acid (2-AA) labelled and purified on Discover DPA-6S SPE columns (SUPELCO). Lipids were eluted in H_2_O and loaded 60:140 H_2_O: MeCN (v/v) for normal phase high-performance liquid chromatography (NP-HPLC). The NP-HPLC system consisted of a Waters Alliance 2695 separations module and an in-line Waters 2475 multi λ-fluorescence detector set at Ex λ 360 nm and Em λ 425 nm. The solid phase used was a 4.6 × 250 mm TSK gel-Amide 80 column (Tosoh Bioscience). A 2AA-labelled glucose homopolymer ladder (Ludger, UK) was included to determine the glucose unit values (GUs) for the HPLC peaks. Individual GSL species were identified by their GU values and quantified by comparison of integrated peak areas with a known amount of 2AA-labelled BioQuant chitotriose standard (Ludger, UK). Results for cell homogenates were normalised to protein content, determined by the bicinchoninic acid (BCA) assay.

GSL species identification and concentration quantification were performed using the software HPLC-RS (https://github.com/ArchercatNEO/HPLC/tree/v0.9).

### Sphingosine analysis by HPLC

Detailed protocol is available at protocols.IO (*36*). Briefly, sphingoid bases were extracted from homogenised cells (0.2 mg protein) in 100 μL H_2_O and spiked with an internal standard (C20, 3 μL 0.03 mM). To the homogenate, 500 μL C:M 1:2 was added and sonicated for 10 minutes, RT. To the samples, 500 μL 1 M NaCl, 500 μL chloroform, and 100 μL 3 M NaOH were added and incubated for 15 minutes, RT, and vortexed every 5 minutes. Samples were then centrifuged (16 000 g/10 minutes) and the lower phase purified on SPE NH2 columns (Biotage) pre-equilibrated with 2 × 1 mL chloroform and eluted with 900 μL acetone. The column elutant was then dried down under N_2_ and resuspended in 50 μL pre-warmed HPLC-grade EtOH. Sphingoid bases were labelled with 50 μL OPA labelling solution (o-phtalaldyhyde dissolved in 20x EtOH, 1× 2-mercaptoethanol and diluted in 2000× 3% boric acid pH 10.5) and incubated in the dark at 20°C for 20 minutes, vortexing at 10 minutes intervals. Samples were buffered in 100 μL MeOH:5 mM Tris pH 7 9:1, centrifuged (800 g / 2 min), and the supernatant was loaded for reverse phase HPLC. Solvent A was MeOH, solvent B was H_2_O, solvent C was MeCN, and solvent D was MeCN:H_2_O 1:4. The RP-HPLC system consisted of a VWR Hitachi Elite LaChrom HPLC system with a L-2485 fluorescence detector set at Ex λ340nm and Em λ455nm. The solid phase used was a Chromolith Performance RP-18e 100–4.6 HPLC column (Merck, Darmstadt, Germany). Results were normalised to protein content.

### Processing of human brain RNA-seq data

FASTQ files from a publicly available dataset on Lewy body diseases (PMID: 34309761), containing samples from the frontal cortex of 7 PD, 6 PDD, 6 DLB and 5 age-matched non-neurological control individuals, was run through a Nextflow (*37*) pipeline that performed quality control (QC) of reads and aligned the samples to the human genome and transcriptome (GRCh38 and Gencode v41 annotations (*38*)). Briefly, initial QC was performed using fastp (v0.23.2 (*39*)), which removed low quality reads and bases, as well as the trimming of adapter sequences. Filtered reads were aligned to the transcriptome using Salmon (correcting for sequence-specific, fragment GC-content and positional biases; v1.9.0 (*40*)) using the entire genome as a decoy sequence. For splicing analyses, reads were aligned to the genome using STAR (v2.7.10a, with two-pass mode). FastQC (v0.11.9 (*41*)) was used to generate read QC data before and after fastp filtering. RSEQC (v4.0.0 (*42*)), Qualimap (v2.2.2a (*43*)) and Picard (v2.27.5 (*44*)) were used to generate alignment QC metrics. MultiQC (v1.13 (*45*)) was used to visualise and collate quality metrics from all pipeline modules. Full details for the pipelines and parameters used for each module can be found in (Brenton (*46*)):

https://github.com/Jbrenton191/RNAseq_splicing_pipeline.

Transcriptome aligned reads generated from Salmon were used to identify covariates. Salmon read files were imported into R (v4.3.0) and converted into gene-level count matrices using the tximport (v1.30.0) and DESeq2 (v1.42.1) packages.

Sample and sequencing covariates were assessed for collinearity using a custom script to reduce the number of covariates tested. This employed the stats (v4.2.3) and Hmisc (v5.2-0) packages to generate Spearman correlations, Kruskal-Wallis and chi-square tests to assess relationships between covariates. Of the numeric covariates that were collinear, those with the lowest overall correlations to all other variables were kept. VariancePartition (v1.32.5) scoring and correlations between remaining covariates and the top 10 principal components (PCs) were then used to assess which covariates should be controlled for in the final analysis.

### DESeq2 differential gene expression

Differential gene expression analysis was performed with DESeq (v1.42.1) using covariates generated as above. Genes with 0 counts in at least one sample in all of the case or control groups, as well as any gene overlapping ENCODE blacklist regions of the genome were removed prior to analysis. Genes were classified as significantly upregulated if their Benjamini and Hochberg FDR adjusted *p* value was ≤ 0.05 and Log2 fold change was > 0, and significantly downregulated if their FDR ≤ 0.05 and Log2 fold change was < 0.

### Single nuclear gene expression analysis of PMCA family and NPC1 interactome

Publicly available processed single nuclear gene expression results from the same samples as used for above bulk gene expression (PMID: 34309761) were downloaded and interrogated for significant changes in genes in the PMCA and NPC1 family of genes and their interaction partners. All quality control, alignment and differential expression analysis measures are described in (PMID: 34309761).

### Differential splicing of bulk RNAseq data from human brain

Leafcutter (0.2.9 (*47*)) was used to perform differential splicing. SJ.out files containing junction reads were first converted to junction files with junctions overlapping ENCODE blacklist regions removed using a custom R script. Junction files were then clustered using the leafcutter_cluster.py script from the leafcutter authors. The following options were used to remove junctions/introns with the following criteria: junctions supported by < 30 reads across all samples, supported by < 0.1% of the total number of junction read counts for the entire cluster and larger than 1 Mb.

Pairwise group comparisons, correcting for identified covariates, were made using Leafcutter’s differential splicing.R function with the following options: intron clusters with more than 40 junctions and a coverage of ≤ 20 reads in ≤ 3 samples in per group were removed, as well as individual introns that were not found in ≥ 5 samples. Clusters were considered significant if they had an FDR-corrected *p* value of ≤ 0.05.

*ATP2B2* splice variant analysis was performed by comparing the delta percent spliced in or delta usage of specific junctions representing splice variants a and b in *ATP2B2* between cases and controls. The coordinates for a variants are start:10329125 and end: 10331983 and start:10329125 and end: 10338176 for b variants.

### Plot generation

Plots were made using ggplot (v3.5.1) and ggtranscript (v0.99.9).

### Conserved Sequence Alignment

Cross-species amino acid and nucleotide alignments were inspected via the UCSC Genome Browser (https://genome.ucsc.edu/cgi-bin/hgTrackUi?g=cons100way) (*48*) using hg19 human and rn6 rat genome assemblies.

### *In vitro* cytosolic calcium measurements using Fura-2

PC12 cells plated on poly-D-lysine coverslips and differentiated to neurons were loaded with the ratiometric fluorescent Ca^2+^ indicator Fura-2 by incubation for 1 h at RT with the acetoxymethyl ester form of Fura-2 (Fura-2 AM, 2.5 µM) and 0.005% (vol/vol) pluronic acid (both from Invitrogen) in HEPES-buffered saline (HBS) containing 10 mM HEPES, 1.25 mM KH_2_PO_4_, 2 mM MgSO_4_, 3 mM KCl, 156 mM NaCl, 2 mM CaCl_2_, and 10 mM glucose (all from Sigma-Aldrich) at pH 7.4. After labelling, cells were washed three times in HBS and mounted in an imaging chamber (Bioscience Tools) with 1 ml HBS.

Epifluorescence images were captured every 3 s with a Megapixel monochrome cooled coupled device camera attached to an Olympus IX73 inverted fluorescence microscope that was fitted with a 20X objective, and a Cairn CoolLED multiple wavelength LED source under the control of MetaFluor 7.10.3.279 software. The Fura-2 ratio was recorded using dual-excitation wavelengths of 340 and 380 nm, and a 440-nm long-pass filter was used to collect emitted fluorescence. GFP-tagged constructs were identified by excitation at 488 nm and capturing emitted fluorescence using a 515 nm long-pass filter.

PC12 neurons were depolarised by replacing HBS with high-potassium buffer, containing 10 mM HEPES, 1.25 mM KH_2_PO_4_, 2 mM MgSO_4_, 156 mM KCl, 3 mM NaCl, 2 mM CaCl_2_, and 10 mM glucose (all from Sigma-Aldrich) at pH 7.4 such that the final KCl concentration was 50 mM. Cells were also acutely challenged with a combination of the TPC2 agonists TPC2-A1N and TPC2-A1P, both at 30 μM.

### *In vitro* cytosolic calcium measurements using Calbryte-590/AM

PC12 cells plated on poly-D-lysine-coated CELview wells and differentiated to neurons were loaded with 2 µM Calbryte-590/AM in an extracellular medium (ECM) at room temperature for 45 mins, followed by a 15-min de-esterification period. For each run, cells were then washed 3x in Ca^2+^-free medium ECM (without added Ca^2+^ and plus 100 µM EGTA), and imaged immediately in the same medium. Chambers were mounted on a Nikon A1R confocal laser-scanning microscope at room temperature, using a 20x objective (1.25-1.50 zoom) and excited at 561 nm (emission centred on 595 nm). An image was acquired every 2 s during stimulation. The fluorescence of each single cell (> 1600 cells per condition) was plotted and analysed using custom-written Magipix software (Dr Ron Jacob, King’s College London). 12 experiments per condition were performed across 4 different days and plotted as the mean of each experiment.

Note: when testing ER Ca2+ release, we intentionally favoured an IP3-coupled agonist over SERCA inhibition (e.g. with thapsigargin) for two reasons: (a) SERCA inhibitors release Ca2+ via a ‘leak’ pathway and not via the relevant physiological Ca2+ release pathway, so thapsigargin would be testing a different pathway; (b) SERCA inhibition releases Ca2+ from the ER very slowly and this allows other Ca2+-removal pathways (such as the PMCA) to buffer and repress the Ca2+ release profile (*49*), whereas rapid Ca2+ release via IP3 is less susceptible.

### AlphaFold3 structure predictions

All protein models were generated using AlphaFold 3 accessed via the AlphaFold Server with default settings (https://alphafoldserver.com/) (*50*). Proteins modelled included human NPC1 (UniProt ID: O15118-1), human PMCA2 (UniProt ID: Q01814-1), yeast Ncr1 (UniProt ID: Q12200-1), and yeast Pmc1 (UniProt ID: P38929-1). Graphical representations of the predicted protein complex structures were prepared using PyMOL (Schrödinger LLC).

### Statistical analysis

To analyse PMCA2 enrichment in the lysosomal fraction following Tagless LysoIP experiments, we performed a one-way Anova analysis, followed by a Tukey’s multiple comparisons test with a 95% confidence interval. Normality of the data was validated using the Kolmogorov-Smirnov test. To evaluate the effects of knocking down the expression of *Atp2b2* in PC12 neurons on Ca^2+^ levels in basal conditions and in response to TPC2 agonists, KCl and ATP, two-tailed unpaired t tests with 95% confidence interval were performed, using the Prism software (version 10.4.1). The same analysis was used to evaluate the effect of knocking down *Atp2b2* on sphingosine and GSLs levels and to compare the gene expression in *Npc1*-null mice to WT mice.

## Acknowledgments

Figure 3 and Supplementary Figure 4 were completely or partially created using Biorender.

## Funding

Aligning Science Across Parkinson’s grant ASAP-000478 (M.E.F.S., J.W.B., D.T.V., J.H., M.R. and F.P.)

The Royal Society Newton International Fellowship NF171121 (M.E.F.S).

Ara Parseghian Medical Research Fund (M.E.F.S and F.P.).

Wellcome Trust Investigator in Science grant 202834/Z/16/2 (F.P.)

Royal Society Wolfson Research Merit Award (F.P).

NPSuisse (M.E.F.S)

Australian NPC Disease Foundation (ANPDF) (M.E.F.S)

Niemann-Pick UK (NPUK) (M.E.F.S)

Niemann-Pick Research Fund (NPRF) (M.E.F.S)

Wellcome Trust grant 218514/Z/19/Z (R.B.)

The Dolby Foundation (J.H.).

## Author contributions

Conceptualization: MEFS and FMP

Investigation: MEFS, RB, JWB, GP, MGP, RR, DTV, DS, YW, EAF, PL, LSP, AM, and LD.

Software: EAF and JWB

Funding acquisition: MEFS, JH and FMP

Supervision and Resources: DGC, EE, DA, CPP, AG, SN, JH, SP, MR and FMP.

Writing – original draft: MEFS, RB and FMP.

Writing – review & editing: all authors.

## Data and materials availability

All data supporting the findings of this study are within the paper or available from the open repository Zenodo with the identifier DOI 10.5281/zenodo.17209110. Code generated on this study for HPLC data analysis (HPLC-RS) is available from: https://github.com/ArchercatNEO/HPLC/tree/v0.10.0 and from Zenodo with DOI: 10.5281/zenodo.18172656. Code used to analyse transcriptomic data is available from: https://github.com/Jbrenton191/RNAseq_splicing_pipeline. Materials are available upon request. The data, code, protocols, and key lab materials used and generated in this study are listed in a Key Resource Table alongside their persistent identifiers at DOI 10.5281/zenodo.17209110.

