## Supplementary Data for "Alternative Splicing Directs PMCA2 to Lysosomes and is Linked to Neurodegeneration"

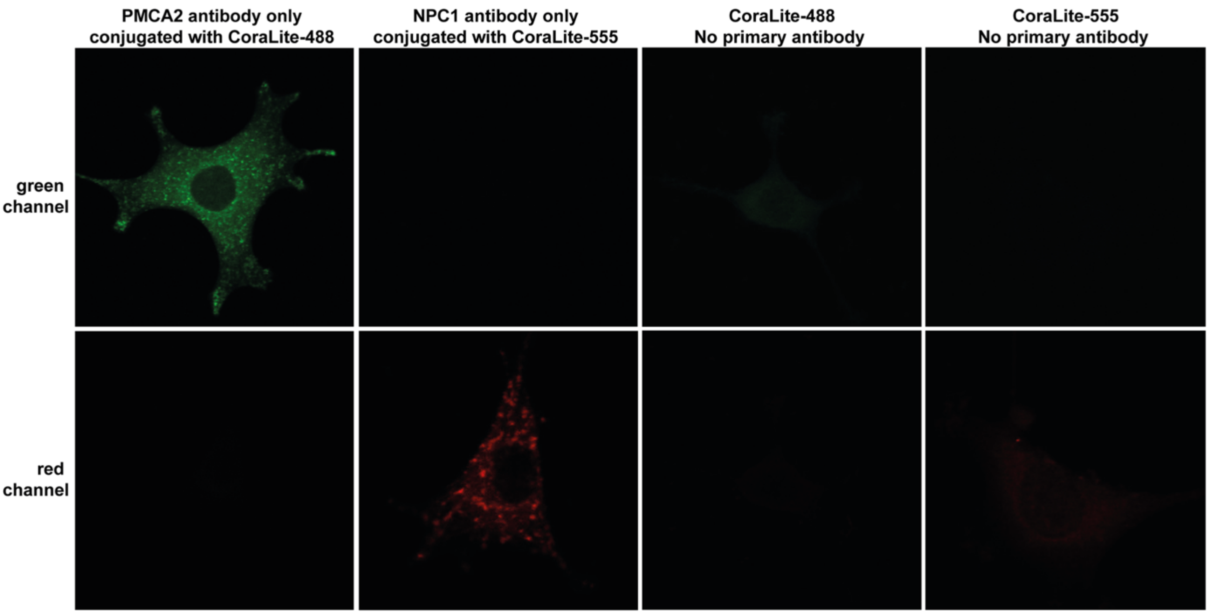


**Supplementary Fig. 1. Immunocytochemistry and confocal microscopy controls in PC12 cells differentiated into neurons.** NPC1 and PMCA2 were detected using primary antibodies directly labelled with CoraLite-555 and CoraLite-488, respectively. No bleed-through was detected when staining with the PMCA2 or NPC1 antibodies only. No significant staining was observed when using CoraLite-555 and CoraLite-488 in the absence of primary antibody. Representative image of four independent experiments.

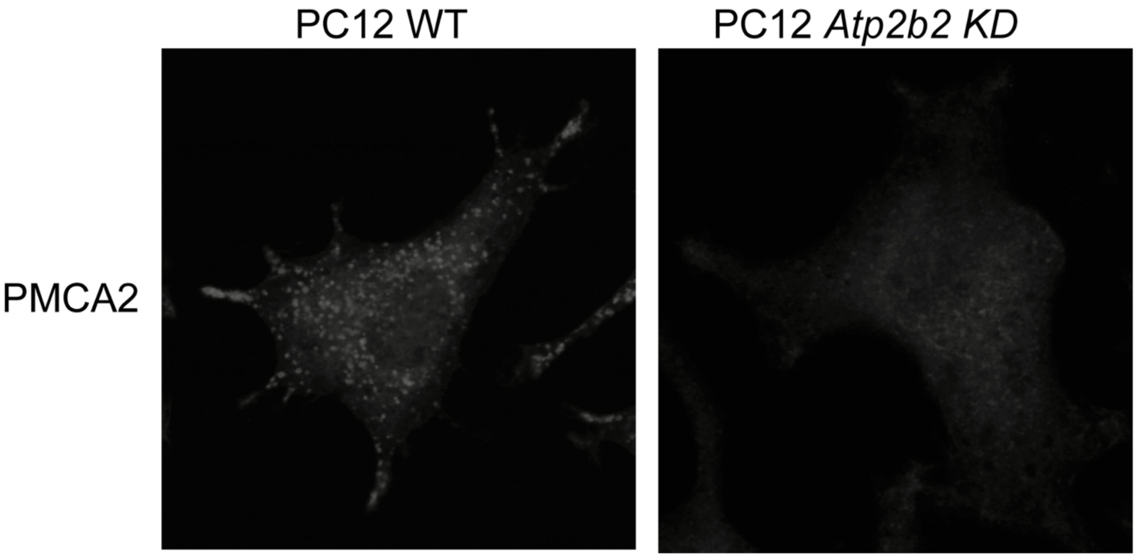

**Supplementary Fig. 2. Polyclonal pan-PMCA2 antibody specificity confirmation in PC12 WT and *Atp2b2* KD cells differentiated into neurons.** PMCA2 was detected using primary antibodies directly labelled with CoraLite-555. A distinct puncta pattern consistent with lysosomal localisation of PMCA2 was detected in wild WT cells, whereas *Atp2b2* KD cells showed a more diffuse background-like staining. Representative image of two independent experiments, 10 fields per experiment.

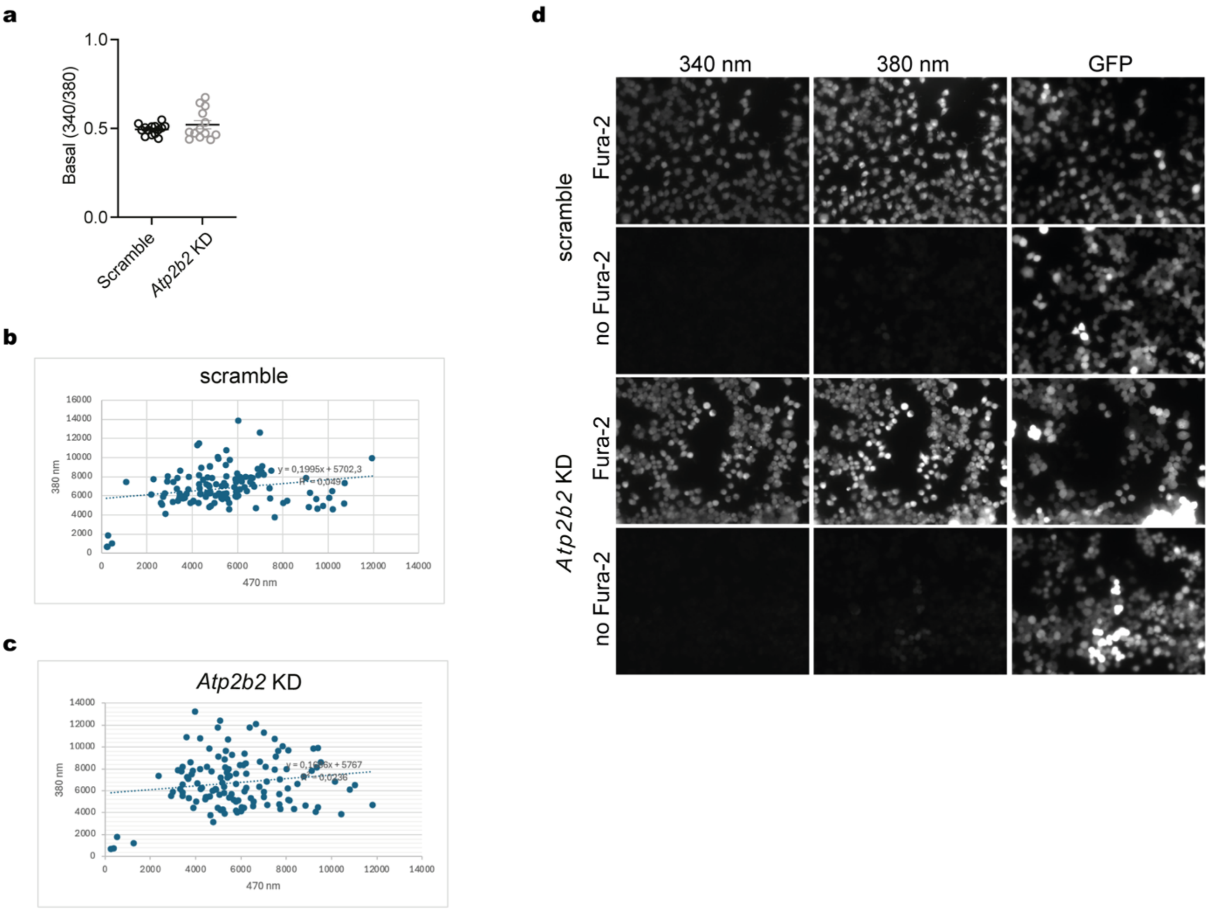


**Supplementary Fig. 3. Basal Fura-2 signal is not affected by *Atp2b2 KD,* and GFP does not interfere with Fura-2 measurements.** PC12 cells stably expressing scramble or *Atp2b2* targeting shRNA were preloaded with the $Ca^{2+}$ probe Fura-2. (a) Basal Fura-2 signal in scramble and *Atp2b2* KD cells. Each point represents the mean response from 30 cells per coverslip from four independent experiments, two to four coverslips per experiment. GFP fluorescence does not leak into the Fura-2 channel in scramble (b) or KD cells (c). (d)

Representative microscopy image showing no correlation between GFP and Fura-2 fluorescence intensities, *n* = 3 independent experiments, 4 coverslips per experiment.

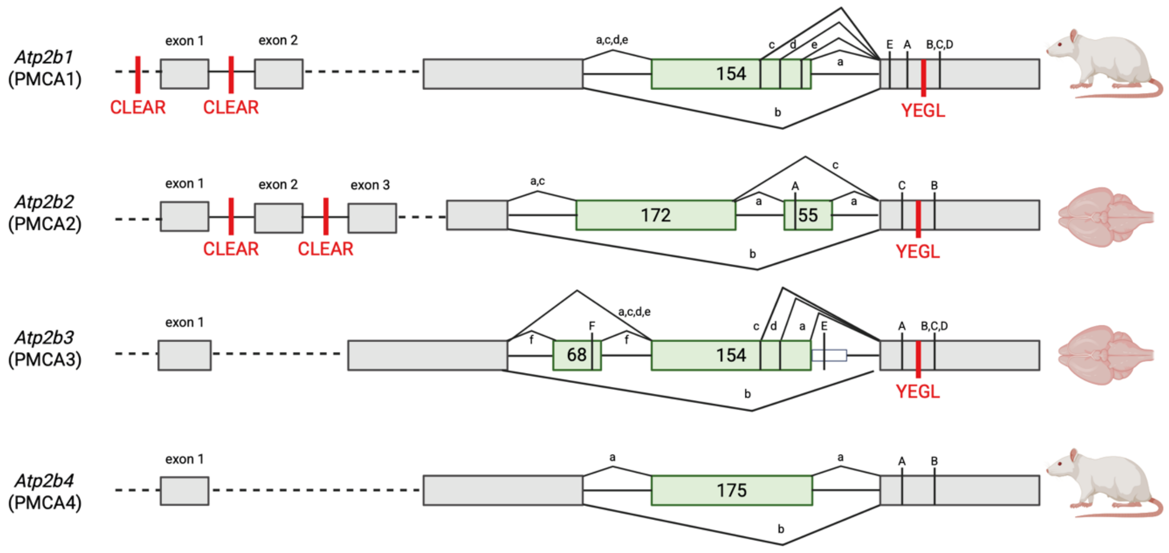


**Supplementary Fig. 4. Specific PMCA splice variants contain the lysosomal targeting motif YEGL**. ***Atp2b1* and *Atp2b2* contain CLEAR sequences.** Alternative splicing in site C of PMCA genes in rat can include or exclude the lysosomal targeting motif YEGL. The exon structure of the different regions affected by alternative splicing is shown for each of the four PMCA genes. Constitutively spliced exons are indicated as grey boxes. Alternatively inserted exons are shown in green. Lowercase letters represent resulting splice variants. Capital letters indicate the position of the translated stop codons. In PMCA3, splice variant "e" results from a read-through of the exon into the following intron (indicated as small white boxes). The sizes of alternatively spliced exons are given as nucleotide numbers. Splice variants shown have supporting sequence data in the literature, RefSeq or Ensembl at the time of submission of this work.

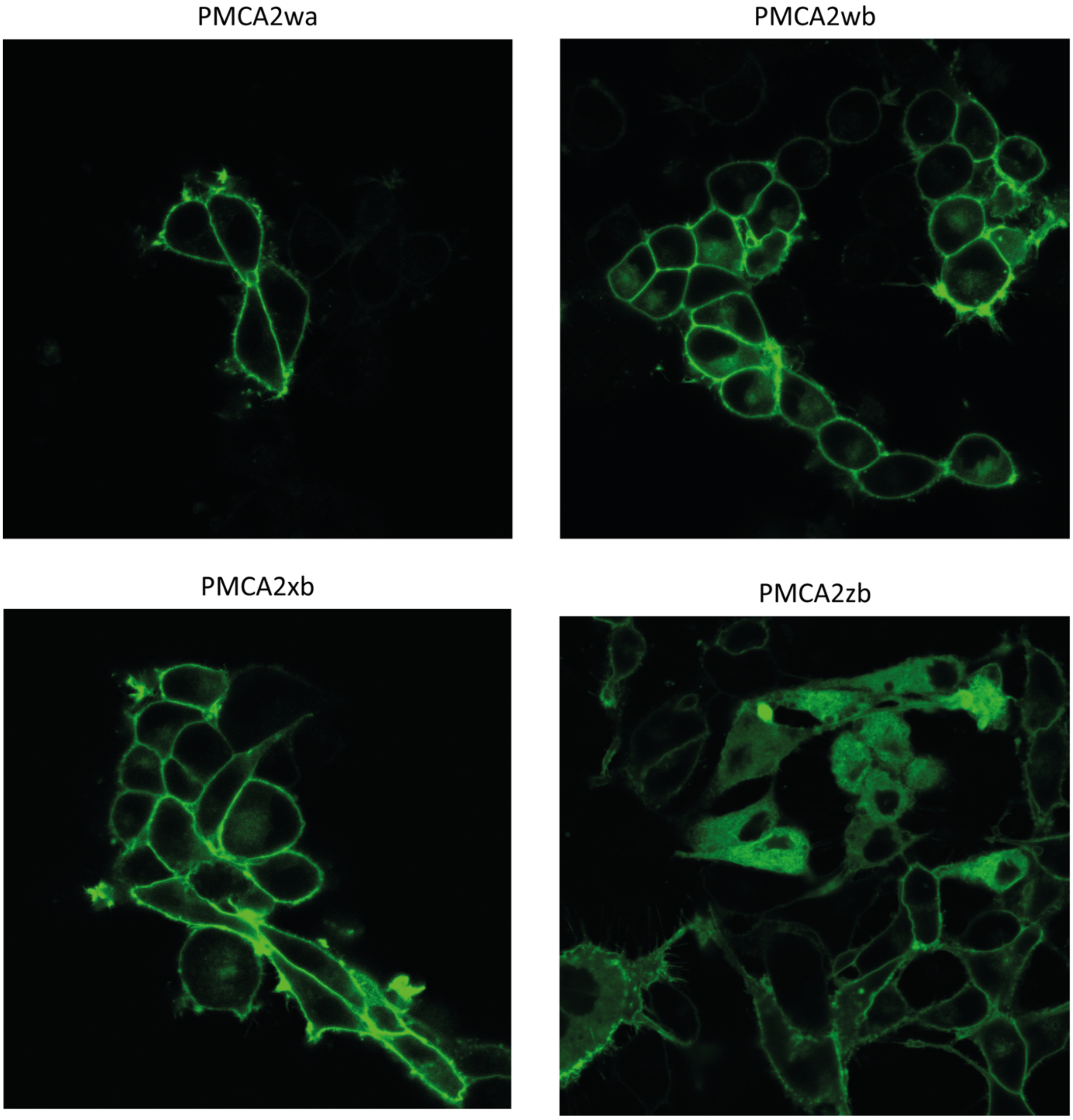


**Supplementary Fig. 5. Subcellular localisation of overexpressed PMCA2 fused to GFP in PC12 cells.** PMCA2wa shows exclusively plasma membrane localisation, while PMCA2wb, PMCA2xb and PMCA2zb exhibit intracellular and plasma membrane localisation when overexpressed. Representative images of 3 independent experiments.

| ΔPSI (%) | | | | |
|---|---|---|---|---|
| | PD | PDD | DLB | |
| splice variant-a | -8.3 | -7.2 | - | |
| splice variant-b | 11.0 | 10.2 | - | *ATP2B1* |
| other | -0.7 | -0.7 | - | |
| splice variant-a | -6.8 | - | 0.9 | |
| splice variant-b | 36.7 | - | -5.7 | *ATP2B2* |
| other | -6.0 | - | 1.2 | |
| splice variant-a | -24.9 | - | - | |
| splice variant-b | 25.7 | - | - | *ATP2B4* |
| other | -0.8 | - | - | |

**Supplementary Table 1. PMCA genes display differential splicing profiles in Lewy body disorders.** Delta percent spliced in PD, PDD and DLB compared to controls for junctions producing alternative splice variants a and b. Only statistically significant changes, FDR significance <0.05, are represented *n* = 5 controls, *n* = 7 PD, *n* = 6 PDD and *n* = 6 DLB. PD: Parkinson's Disease, PDD: PD with dementia, DLB: dementia with Lewy bodies.

| Gene | Forward | Reverse | use | splice variant amplified | amplicon size |
|---|---|---|---|---|---|
| | | | | a | 552 bp |
| *Atp2b2* | CACCATCCCTACCAGCAGGC | CAGGTCGGTGTCATCGATGA | PCR | b | 325 bp |
| | | | | c | 497 bp |
| | GCTACAGGACGTGACACTTATC | CCCCGGAACTGTTTCTCCTTA | qPCR | total | 237 bp |
| | ATCCAATGCTCTTTCCTCTCCG | TTCACGACGCGGATCTCTTTA | qPCR | a | 119 bp |
| *Atp2b2* | AGGATGTGGAAGAGATAGACCAC | ACGACGCGGATCTGTGTC | qPCR | b | 100 bp |
| | AGCCAGAGCCAGGACGTAG | CTTCACGACGCGGATGCCC | qPCR | c | 115 bp |
| *Rpl37* | CATCCTTTGGTAAGCGTCGCA | TGGCACTCCAGTTATACTTCCT | qPCR | - | 138 bp |

**Supplementary Table 2. Mouse primers used in PCR experiments.**
